# Anywhere but here: local conditions alone drive dispersal in *Daphnia*

**DOI:** 10.1101/334243

**Authors:** Philip Erm, Matthew D. Hall, Ben L. Phillips

## Abstract

Dispersal is fundamental to population dynamics and it is increasingly apparent that, despite most models treating dispersal as a constant, many organisms make dispersal decisions based upon information gathered from the environment. Ideally, organisms would make fully informed decisions, with knowledge of both intra-patch conditions (conditions in their current location) and extra-patch conditions (conditions in alternative locations). Acquiring information is energetically costly however, and extra-patch information will typically be costlier to obtain than intra-patch information. As a consequence, theory suggests that organisms will often make partially informed dispersal decisions, utilising intra-patch information only. We test this proposition in an experimental two-patch system using populations of the aquatic crustacean, *Daphnia carinata*. We manipulated conditions (food availability) in the population’s home patch, and in its alternative patch. We found that *D. carinata* made use of intra-patch information (resource limitation in the home patch induced a ten-fold increase in dispersal probability) but made no use of extra-patch information (resource limitation in the alternative patch did not affect dispersal probability). Our work highlights the very large influence that information can have on dispersal probability, but also that dispersal decisions will often be made in only a partially informed manner. The magnitude of the response we observed also adds to the growing chorus that condition-dependence may be a significant driver of variation in dispersal.

## Introduction

It is increasingly appreciated that dispersal is, like survival and reproduction, a fundamental facet of life history (Bonte and Dahirel 2017). Dispersal behaviours can have profound consequences for individual fitness, and the variety of dispersal behaviours in a population not only influences local population dynamics, but also metapopulation and evolutionary dynamics across a species range (Clobert et al. 2001). For reasons of simplicity, the majority of spatially-explicit ecological models assume that dispersal is both uninformed and unchanging; that individuals disperse at fixed rates, and that they do so without recourse to information about environmental conditions (*e.g.*, by default in models utilising reaction-diffusion or integrodifference equations; Fisher 1937; Skellam 1951; Kot et al. 1996; but see Fronhofer et al. 2016). There is now considerable evidence, however, that dispersal decisions are routinely informed by aspects of the environment (Clobert et al. 2009), and such information use can be expected to have non-negligible effects on ensuing population and evolutionary dynamics (Delgado et al. 2014; Ponchon et al. 2015; Urban et al. 2016).

The most common form of informed (or condition-dependent) dispersal is density-dependent dispersal (Bowler and Benton 2005). Here, individuals acquire information about population density, and, if conditions more favourable to survival and reproduction are likely to be found elsewhere, make the decision to disperse. When all else is equal, high density – with its greater competition for resources, greater rates of disease transmission and so on – will be associated with poorer conditions (Bowler and Benton 2005). It has been repeatedly demonstrated that individuals acquire information on density and act upon it (De Meester and Bonte 2010; Fellous et al. 2012; Martorell and Martínez-López 2014; Fronhofer et al. 2017a). In many species, this information is acquired through food limitation (Fronhofer et al. 2017b): when resources are limited, particularly in juvenile stages, individuals tend to be more dispersive. Information on the relative merits of different alternative locations can also be acquired in numerous ways, including prospecting (actively moving through, and assessing, new locations, *e.g.*, Pärt and Doligez 2003) and by observing immigrating conspecifics (Cote and Clobert 2007).

Several models have now been constructed to examine the evolution of informed dispersal (*e.g.*, Bocedi et al. 2012; Delgado et al. 2014) and, while they are in general agreement that informed dispersal will often evolve, much hinges on the ease with which information is acquired, along with its value (Poethke et al. 2016). In an ideal world, we would expect dispersal decisions to be made on a balance of ‘push’ factors, such as local patch conditions, and ‘pull’ factors, such as the quality of other patches. The quality of the new patch should be high enough (relative to the home patch) that it offsets the fitness costs of moving. But organisms do not live in an ideal world: information can be costly in terms of time and energy to acquire (Bonte et al. 2012), and there is a likely asymmetry of cost such that information about alternative patches is harder to obtain than information about an individual’s home patch. Thus, it may often be the case that individuals act on the limited information that is most easily acquired: intra-patch information.

Still, broader ecological and population dynamics may be powerfully influenced by the relative strength of the pull exerted by extra-patch information. In metapopulations with dispersers that can take advantage of extra-patch information, patch persistence may increase if patches with perilously low population sizes – but abundant resources as a result – become more appealing to dispersers. Indeed, simply being able to detect other conspecifics in these vulnerable patches may have the same effect (Clobert et al. 2009), although patches with suitable habitat may conversely become overpopulated if they attract a disproportionate number of migrants. In biological invasions, invasion speed may be boosted if colonisers are able to use extra-patch information to select suitable habitats, or hindered if it instead causes them to favour ecological traps (Kokko 2006). If intra-patch information is dominant in motivating dispersal however, invaders may instead be expected to distribute themselves indiscriminately, rendering the invasion highly sensitive to both the proportion of suitable habitat in the landscape and any temporal fluctuations in its quality (Neubert et al. 2000; Schreiber and Lloyd-Smith 2009). In organisms generally then, gauging the relative strength of the push caused by intra-patch information and the pull caused by extra-patch information will help to resolve questions of such a character.

Here, we examine their relative influence on dispersal by manipulating food resource levels in experimental populations of the aquatic crustacean, *Daphnia carinata*. Dispersal in *Daphnia* is usually characterised as being driven by the passive transport of ephippia (long-lived resting eggs) by water fowl or other vectors (Allen 2007; Frisch et al. 2007; Van de Meutter et al. 2008); however, individuals can also actively disperse between permanently or temporarily interconnected water bodies (Michels et al. 2001; Cottenie et al. 2003). Although it has been demonstrated that *Daphnia* do boost ephippia production in response to information cues indicating low local resource availability (Carvalho and Hughes 1983; Hobaek Anders and Larsson Peter 1990; Kleiven et al. 1992), a greater range of responses have been observed regarding its effects on active movement. Environments with relatively higher food concentrations have been shown to increase *Daphnia* movement behaviours like swimming speed and sinking rate (Dodson et al. 1997); however, in other instances, they have been shown to slow movement, with much depending on the *Daphnia* species or clone line under examination (Young and Getty 1987; Larsson and Kleiven 1997; Roozen and Lürling 2001). *Daphnia* have also been seen to adhere to ideal free distributions under ordinary circumstances, with individuals favouring regions of high food concentration so long as they fall within natural ranges (Jakobsen and Johnsen 1987; Neary et al. 1994; Jensen et al. 2001). It would appear likely then, that *Daphnia* exploit information to regulate their dispersal efforts between patches. It is less clear however, if this behaviour is governed entirely by intra-patch information, or if extra-patch information also influences dispersal propensity.

Using *D. carinata*, we determine if individuals respond to intra-patch resource levels by modifying their rates of active dispersal between linked patches in small multi-patch mesocosms. We also ask whether this response is contingent upon extra-patch conditions; the presence or absence of *ad libitum* food in the neighbouring patch.

## Methods

### Laboratory population of *D. carinata*

All *D. carinata* used were genetically identical members of a single clone line. The founding member of this lineage was collected at 38°10’34.3“S, 144°21’14.1”E (a lake in Geelong, Victoria, Australia) in October 2016. Its offspring were used to establish laboratory stock cultures, which were housed in glass jars containing 300 ml of ADaM zooplankton medium (according to the recipe of Klüttgen et al. 1994; as modified by Ebert 2013) and kept within growth chambers maintained at 22° C on a 12.30 light:11.30 dark photoperiod. Stocks were fed the green algae *Scenedesmus*. In order to reduce any potential impact of maternal effects, all individuals used in the experiment were taken from stocks that were maintained under these conditions for at least two generations.

### Experimental materials and conditions

We set up two-patch microcosms within which to measure dispersal of *Daphnia*. Each patch was a 950 ml plastic Cryovac XOP-0980 container filled with 600 ml of ADaM and kept on bench tops in an open air laboratory. The laboratory was maintained at 22° C and each container was covered with a transparent plastic sheet that was only removed during feeding and data collection. From their base, each container’s dimensions were 90 mm x 75 mm x 110 mm, and they possessed a gradual taper from top to bottom, such that the opening at the top of the container (100 mm x 90 mm) was larger than the base of the container. A circular hole with a diameter of 15 mm was centrally located 35 mm above the base on one of the long sides of each container. This was connected to plastic PVC piping of an identical internal diameter that linked one container to the next, acting as a 117 mm long tunnel through which *D. carinata* could disperse between the two containers. At the commencement of the experimental trials, dispersal between containers was prevented by inserting cotton balls into the openings of the connecting tunnel.

### Food availability experiment

Within this two-patch system, we examined the effect of intra- and extra-patch food availability on the dispersal rate of *D. carinata*. We seeded one half of each two-patch system with 10 adult females taken from stock cultures, and allowed this population to grow for 9 days in the experimental system while dispersal was blocked. This resulted in an average of 122 individuals across a variety of age and size classes in each replicate when dispersal commenced. On the 10th day, we then unblocked the dispersal tunnels and counted the number of individuals in each patch every 24 hours thereafter for four days.

Our patch pairs were allocated to four treatment combinations (n = 5 per combination) according to a two factor crossed design in which we independently modified food availability in the two patches. Factor 1 was intra-patch food availability: once the dispersal tunnel was unblocked, half of the populations no longer received food in their starting patch. Factor 2 was extra-patch food availability: here we either daily added food to the second patch (commencing on day 7, three days before dispersal was allowed) or withheld food altogether from this patch. This meant that half of the populations were dispersing into patches that contained no food at all, and the other half into patches with an abundance of food. Food in this case was a daily fed mixture of 8 million *Scenedesmus* sp. cells (an unidentified Australian *Scenedesmus*) and 12 million *Scendesmus obliquus* cells.

We examined the effect of feeding regimes on absolute population sizes using ANOVA. Here, we examined two response variables: the total population size at 96 hours (summed across both patches); and the population size in patch 2 at 96 hours. We compared the proportion of individuals that had reached patch 2 after 96 hours between treatment combinations using a generalised linear model with binomial errors and a logit link, with each individual in each patch being characterised as a trial in which either success (dispersing into patch 2) or failure (remaining in patch 1) had resulted. All statistical tests were performed in R version 3.5.0 (R Core Team 2018). All experimental data is available in the figshare repository at https://doi.org/10.4225/49/5b0f62dc23b4c.

## Results

Over the course of the dispersal phase, total population sizes across both patches generally increased or decreased according to whether patch 1 was fed or not, with fed treatments overall growing in size and unfed treatments shrinking (Figure 2). At 96 hours, we found a significant effect of food availability in patch 1 on total population size across both patches (*F*_1,16_ = 10.826, *P* < 0.01; Table 2), but not of food availability in patch 2 (*F*_1,16_ = 0.013, *P* = 0.912; Table 2). The interaction between feeding treatments in the two patches was also not significant with regard to total population size (*F*_1,16_ = 0.481, *P* = 0.498; Table 2).

Examining the proportion of individuals dispersing at 96 hours, we found no significant effect of the interaction between intra- and extra-patch feeding treatments (*z* = 1.073, *P* = 0.283; Table 1), and likewise no significant effect of food availability in patch 2 (*z* = 0.138, *P* = 0.890; Table 1). We did however find a significant effect of food availability in patch 1 (*z* = 10.843, *P* < 0.001; Table 1), with intra-patch food deprivation resulting in an approximately ten-fold higher proportion of the total population dispersing (food-deprived patch 1, mean = 0.259, SE = 0.0374; well-fed patch 1, mean = 0.0218, SE = 0.00718; Figure 1).

**Table 1:** Statistical test results for differences in the proportion of *D. carinata* dispersing depending on food availability in patch 1 and food availability in patch 2. A generalised linear model was used with parameter estimates on the logit scale and binomial errors. Model variance was checked for overdispersion and did not violate standard GLM assumptions.

| Parameter | Estimate (SE) | *z* stat | *P* value |
| --- | --- | --- | --- |
| *Intercept* | -0.938 (0.100) | 9.371 | <0.001 |
| *Food available in patch 1* | -2.872 (0.265) | 10.84 | <0.001 |
| *Food available in patch 2* | -0.0206 (0.150) | 0.138 | 0.890 |
| *Food available in patch 1* × *Food available in patch 2* | 0.375 (0.350) | 1.073 | 0.283 |

**Figure 1:**
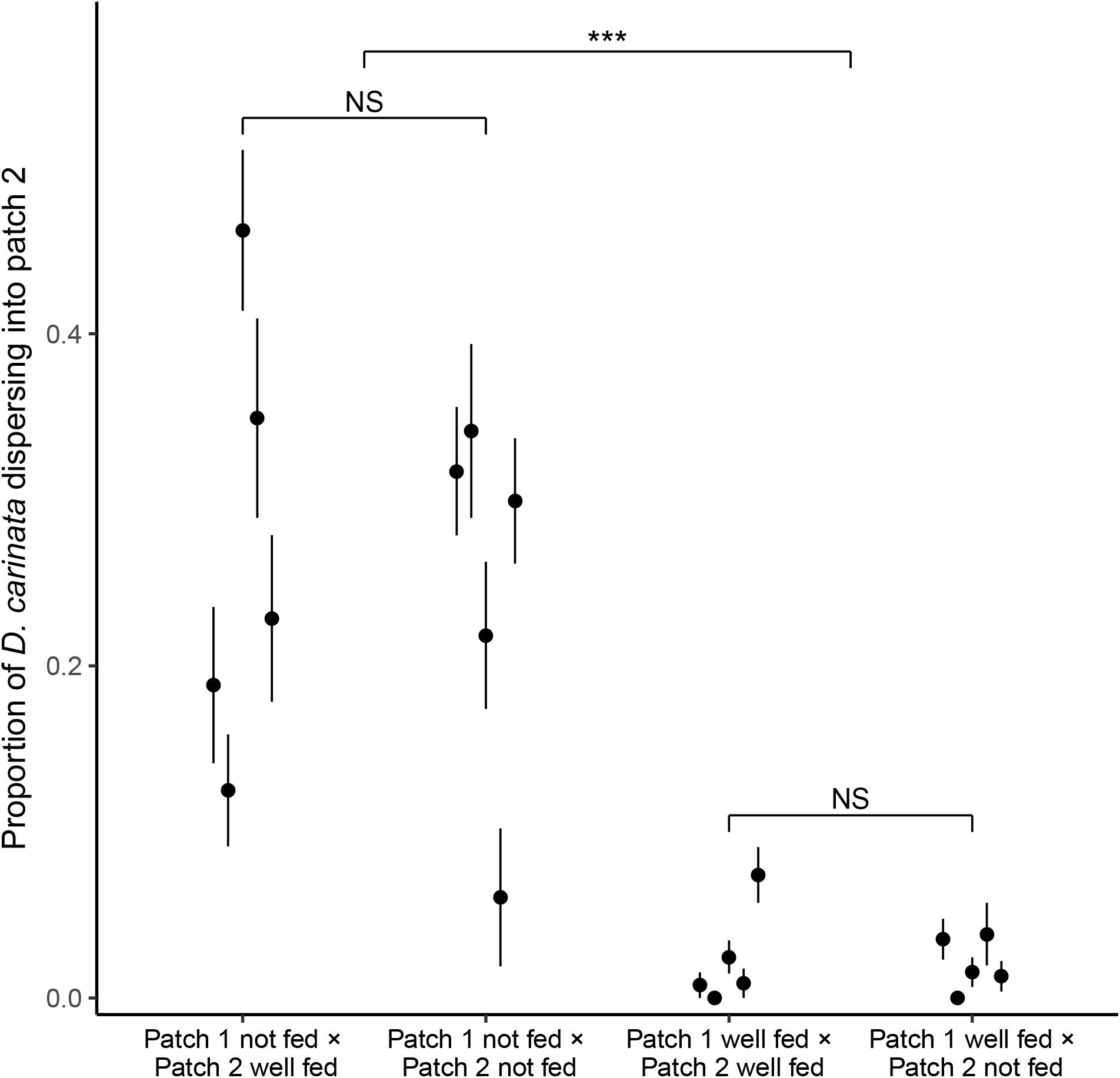
The effect of 96 hours of food deprivation on the proportion of *D. carinata* that had dispersed into patch 2, both with and without food available in patch 2 (n = 5 container pairs per treatment combination). Each point and line is given by the mean number of *D. carinata* individuals in patch 2 as a proportion of the total population size between the two patches ±SE (see Table A1 for precise results). Food availability in patch 1 alone was found to have a significant effect on the proportion of the population that had dispersed (*P* < 0.001).

An analysis based on absolute numbers in patch 2, rather than proportions, yielded the same overall pattern. Here, we found a significant difference in the total number of individuals in patch 2 according to whether patch 1 had been fed or not (*F*_1,16_ = 13.605, *P* < 0.01; Table 2), but no significant effect of food availability in patch 2 (*F*_1,16_ = 0.102, *P* = 0.754; Table 2). Indeed, patch 1 unfed groups had a far higher number of individuals in patch 2 despite their significantly lower total population sizes (individuals in patch 2: food-deprived patch 1, mean = 25.1, SE = 5.12; well-fed patch 1, mean = 4.3, SE = 1.73).

**Table 2:** ANOVA test results for differences in the total population sizes and absolute number of dispersers depending on food availability in patch 1 and food availability in patch 2 for *D. carinata*. Standard ANOVA assumptions were not violated.

| Source | *df* | *F* stat | *P* value |
| --- | --- | --- | --- |
| Total population sizes |  |  |  |
| *Food available in patch 1* | 1 | 10.826 | <0.01 |
| *Food available in patch 2* | 1 | 0.013 | 0.912 |
| *Food available in patch 1* × *Food available in patch 2* | 1 | 0.481 | 0.498 |
|  | 16 |  |  |
| Absolute dispersers |  |  |  |
| *Food available in patch 1* | 1 | 13.605 | <0.01 |
| *Food available in patch 2* | 1 | 0.102 | 0.754 |
| *Food available in patch 1* × *Food available in patch 2* | 1 | 0.408 | 0.532 |
|  | 16 |  |  |

## Discussion

In our system, there was a significant increase in inter-patch dispersal when *D. carinata* were deprived of food (Figure 1), indicating that *D. carinata* exploited intra-patch information to inform their dispersal decisions. By contrast, extra-patch conditions (food either abundant or entirely absent) had no effect on dispersal rates. Thus it appears that *D. carinata* either did not use, or were incapable of using, extra-patch information to inform their dispersal decisions.

Our first result – that animals increase dispersal propensity when faced with local resource shortages – has been well established empirically. Studies on taxa ranging from plants to invertebrates and vertebrates either imply, or experimentally demonstrate, that resource shortage is a powerful piece of information motivating dispersal (*e.g.*, Bowler and Benton 2005; Martorell and Martínez-López 2014; Fronhofer et al. 2017b). Our study adds *D. carinata* to the long list of organisms that exploit this piece of intra-patch information. Due to the generality of this phenomenon, it also appears likely that comparable results would be seen for other species of *Daphnia*.

That the dispersal we observed was indeed driven conditionally by resource shortage, rather than density in and of itself, becomes obvious when examining patch 1 population sizes across treatments. Since the nature of our experimental design precluded any attempt to control density, those treatments that were well fed in patch 1 kept growing over time compared to those that were not, manifesting in a significantly higher density level by the end of the experiment (Figure 2). Despite this advantage in density, which would not only have created more potential dispersers but also exacerbated any density-driven push effect, a significantly greater number of individuals dispersed in the treatments experiencing lower densities. Density, independent of resource shortage, has been demonstrated to cause changes in life-history in *Daphnia* spp. (Matveev 1993; Burns 1995, 2000); but here, where dispersal is considered, intra-patch resource shortage appears to be a far more powerful driver than density per se.

**Figure 2:**
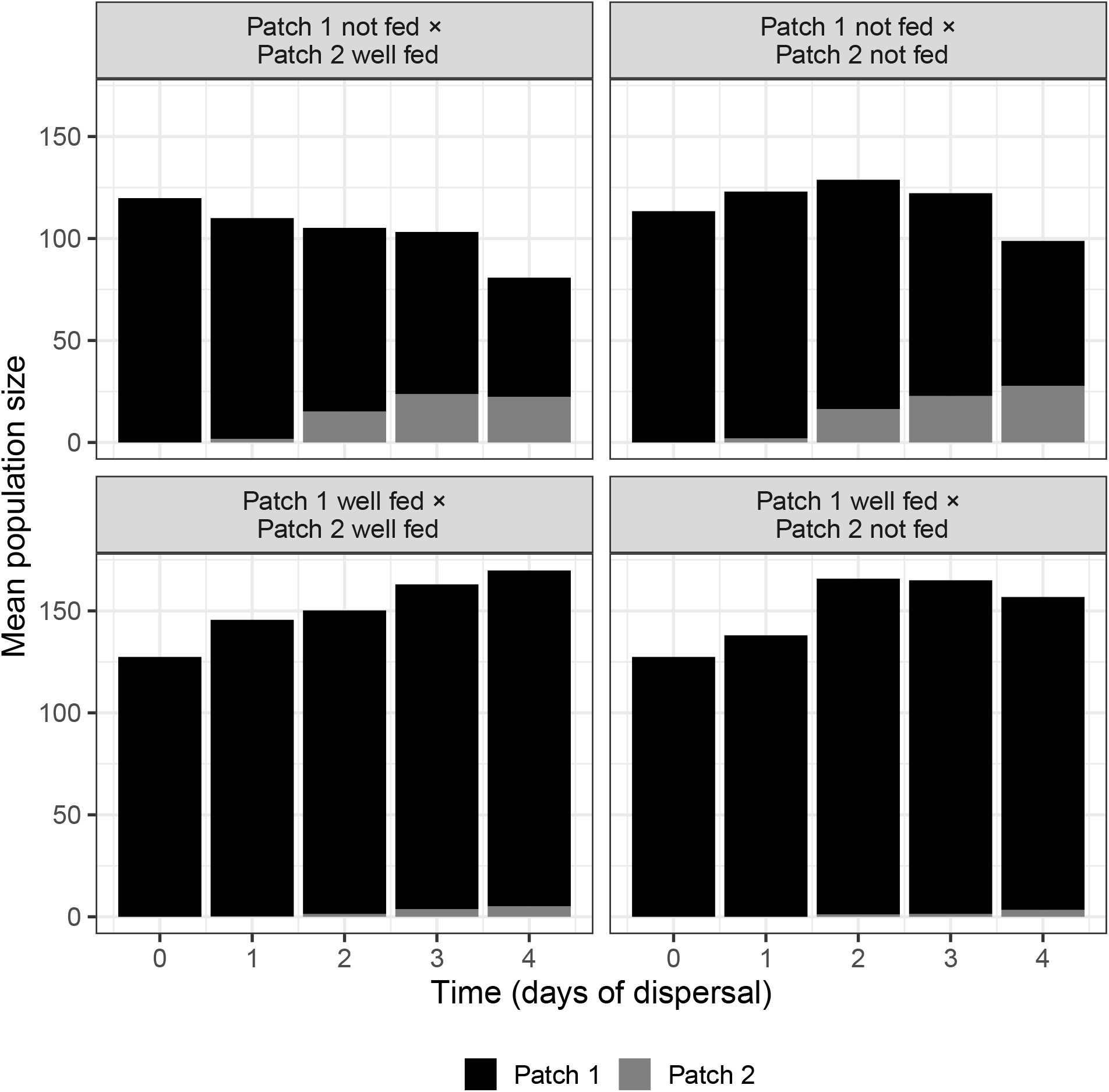
The effect of feeding regime on mean population size across both patch 1 and patch 2 over the dispersal phase (n = 5 container pairs per treatment combination). Bars are stacked, such that both patch 1 and patch 2 population sizes combine to indicate mean population size.

The magnitude of the dispersal increase we observed also indicates that the effect of local resource information on dispersal rates may be extremely pronounced. In terms of the proportion of dispersers, 25.9% of individuals dispersed into patch 2 under food deprivation, whereas less than a tenth of that (2.18%) did so under well-fed conditions (Figure 1). Although this particular measure may have been inflated by the population growth that continued to occur in the well-fed treatments, the large difference in the absolute number of dispersers (food-deprived patch 1, 25.1 dispersers; well-fed patch 1, 4.3 dispersers) despite the afore-mentioned higher density in the well-fed treatments reiterated the strength of the effect. This suggests that ecological models may benefit substantially by accounting for conditional factors, like resource availability, that may have a very large effect on dispersal behaviour.

Our second result – that favourability of conditions in the second patch had no effect on dispersal – highlights the relative importance of push versus pull factors in driving a population’s movement. In the present case, to obtain information that would draw *D. carinata* into the second patch, individuals either had to engage in prospecting within the inter-patch tunnel and the second patch, or to sense extra-patch conditions remotely. We found no evidence to suggest that either source of information was exploited. In terms of more direct means of gathering information, extra-patch information gathering behaviours like prospecting are predicted to be costly due to the threat of predation that comes from moving into novel environments (Bonnet et al. 1999; Hiddink et al. 2002; Bonte et al. 2012), or the simple energetic cost of having to move to assess new patches (Delgado et al. 2014). In *D. carinata* specifically, it seemed much more likely that chemoreception would serve as the primary means of ascertaining extra-patch conditions, as chemical signals from both conspecifics and other organisms have been demonstrated to have a multitude of effects on *Daphnia* growth and behaviour (Larsson and Dodson 1993; Dodson et al. 1994). Indeed, it has been previously shown that *Daphnia magna* and *Daphnia pulex* are unaffected by olfactory cues from algae (Roozen and Lürling 2001), but that a *Daphnia galeata* and *Daphnia hyalina* hybrid responds to them (van Gool and Ringelberg 1996). Here however, the dominance of resource limitation in pushing dispersal from the local patch indicated that the pull to move into new patches was relatively weak in comparison, either because obtaining more information was costly, or because that information was in some way imperceptible or ignored.

In conclusion, our results add to the growing body of evidence that condition-dependent dispersal is the norm amongst taxa, and that it is moreover capable of generating substantial differences in dispersal behaviour (Legrand et al. 2015; Fronhofer et al. 2017b). This growing empirical consensus warns against the simplifying assumption – used in the majority of ecological and evolutionary models – that dispersal rate is constant with respect to conditions. Relaxing that assumption is now well justified on empirical grounds, and the magnitude of shift in dispersal resulting from condition dependence suggests that it will have non-trivial effects when incorporated into mechanistic models of evolution, population dynamics, invasion spread, and so on (*e.g.*, Armsworth 2008; Armsworth and Roughgarden 2008). In this light, the relative use of extra- vs intra-patch information is important because when we move to a conditional dispersal model, the obvious simplifying assumption is that organisms exploit only intra-patch information. Our results suggest that intra-patch information is dominant in *D. carinata*, but the degree to which this is true generally will determine how complex our models of dispersal really need to be.

## Acknowledgements

We thank Katrina-Lee Ware and Hannah Edwards for their invaluable assistance in the preparation of the experimental set-up. We also thank two anonymous reviewers for their comments on the manuscript. Funding was provided by the Australian Research Council (DP160101730; FT160100198).

